# Acute session of three endurance exercise intensities alters subcutaneous adipose tissue transcriptome in regular exercisers

**DOI:** 10.1101/2025.05.02.651890

**Authors:** Cheehoon Ahn, Tao Zhang, Thomas Rode, Gayoung Yang, Olivia K. Chugh, Sierra Ellis, Sophia Ghayur, Shriya Mehta, Ryan Salzman, Hui Jiang, Stephen C.J. Parker, Charles F. Burant, Jeffrey F. Horowitz

## Abstract

The primary aim of this study was to compare the acute effects of three exercise intensities on abdominal subcutaneous adipose tissue (aSAT) transcriptome in regular exercisers. A total of 45 adults who exercise regularly were assigned to perform a single session of either low-intensity continuous (LOW; 60min at 30% VO_2_max; n=15), moderate-intensity continuous (MOD; 45min at 65% VO_2_max; n=15), or high-intensity interval exercise (HIGH; 10×1min at 90% VO_2_max interspersed with 1min active recovery; n=15). aSAT biopsy samples were collected before and 1.5hours after the exercise session for bulk RNA sequencing and targeted protein immunoassays. HIGH upregulated genes involved in cytokine secretion, insulin signaling, and proteolysis while MOD and LOW upregulated genes regulating ECM remodeling, ribosome biogenesis, and oxidative phosphorylation pathways. Exercise-induced changes in aSAT angiogenic, MAPK cascade, and clock genes, ERK protein phosphorylation, and circulating cytokines were similar after all three exercise treatments. Network analysis identified exercise-responsive gene clusters linked to cardiometabolic health traits. Cell-type analysis highlighted a heterogeneous response of aSAT cell types to exercise, with distinct patterns observed across exercise intensities. Collectively, our data characterizes early responses in aSAT after a single session of exercise. Because adaptations to exercise training stem from an accrual of responses after each session of exercise, these early responses to exercise are likely important contributors to the long-term structural and functional changes that occur in adipose tissue in response to exercise training.

## INTRODUCTION

The cardiometabolic health benefits of regular exercise are well-documented (1, 2); however, the precise mechanisms driving these effects remain incompletely understood. Most studies examining the metabolic benefits of exercise largely focus on adaptations in skeletal muscle, but far less is known about exercise-induced adaptations in adipose tissue that also underlie some of the health benefits of exercise (3, 4). Exercise training has been found to increase abdominal subcutaneous adipose tissue (aSAT) capillarization and mitochondrial proteins, remodel extracellular matrix (ECM), and lower pro-inflammatory macrophage infiltration (3, 4). However, how exercise triggers these adaptive responses in aSAT, and whether the exercise intensity and/or energy expended during exercise may impact these triggers is unclear.

Adaptations to regular exercise typically result from the accrual of repeated exposure to acute transcriptional changes that occur shortly after each exercise session (5). For example, the increase in skeletal muscle mitochondrial density in response to weeks or months of endurance training is largely due to the repetitive transient increase in transcriptional regulators of mitochondrial biogenesis and transcripts involved in oxidative phosphorylation that occur in the hours after each exercise training session (6–8). The increase in aSAT mitochondrial density that has also been found with exercise training (9, 10), may occur through upregulation of the similar transcriptional pathways as observed in skeletal muscle (11, 12). Additionally, we previously observed a robust increase in the mRNA expression of the key angiogenic transcription factor VEGFA one hour after aerobic exercise (13), which might be part of the initial step for enhanced vascularization in aSAT observed with exercise training (3). Hence, identifying transcriptomic signals after a session of exercise is critical to understanding the mechanisms underlying the adaptations in aSAT that occur with exercise training. Importantly, many of the well-described adaptations to exercise training vary depending on the intensity, duration, and/or energy expended during the exercise training sessions. This is also true for at least some of the adaptations in adipose tissue. For example, it has been reported that high-intensity training induced some more pronounced adaptations in adipose tissue (e.g., greater abundance of capillaries and smaller adipocytes and less pro-inflammatory macrophage infiltration) compared with moderate-intensity training (14, 15).

The primary aim of this study was to compare the acute effects of exercise performed at low-, moderate-, and high-intensity that are commonly prescribed for health and fitness, on aSAT transcriptome. We hypothesized that 1) a single session of exercise would alter aSAT transcriptome, cytokine production, and protein phosphorylation and 2) the exercise-induced effects would be distinct among three different exercise sessions.

## METHODS

### Subjects

We only recruited subjects who regularly engage in endurance exercise to avoid the confounding influence of the stress response of an exercise stimulus in persons who are not accustomed to exercise. Out of 190 subjects who expressed interest in the study, a total of 45 healthy subjects [body mass index (BMI) 20-30kg/m^2^, age 18-40 years] were enrolled and completed the study (Figure 1A). All participants were regular exercisers (≥30min, ≥3 days/week, moderate to vigorous intensity endurance-type exercise) for at least 2 months and reported having stable body weight for at least 6 months. Subjects were not taking any medications or supplements known to affect their metabolism except for contraceptive medications for some female participants. Subjects also had no history of cardiovascular or metabolic disease. All female participants were premenopausal and not pregnant or lactating. All subjects completed a detailed medical history survey and resting electrocardiogram, which were reviewed by a physician before any testing. Written informed consent was obtained from all subjects before the study. This study conformed to the standards set by the Declaration of Helsinki, except for registration in a database, and was approved by the University of Michigan Institutional Review Board.

**Figure 1.**
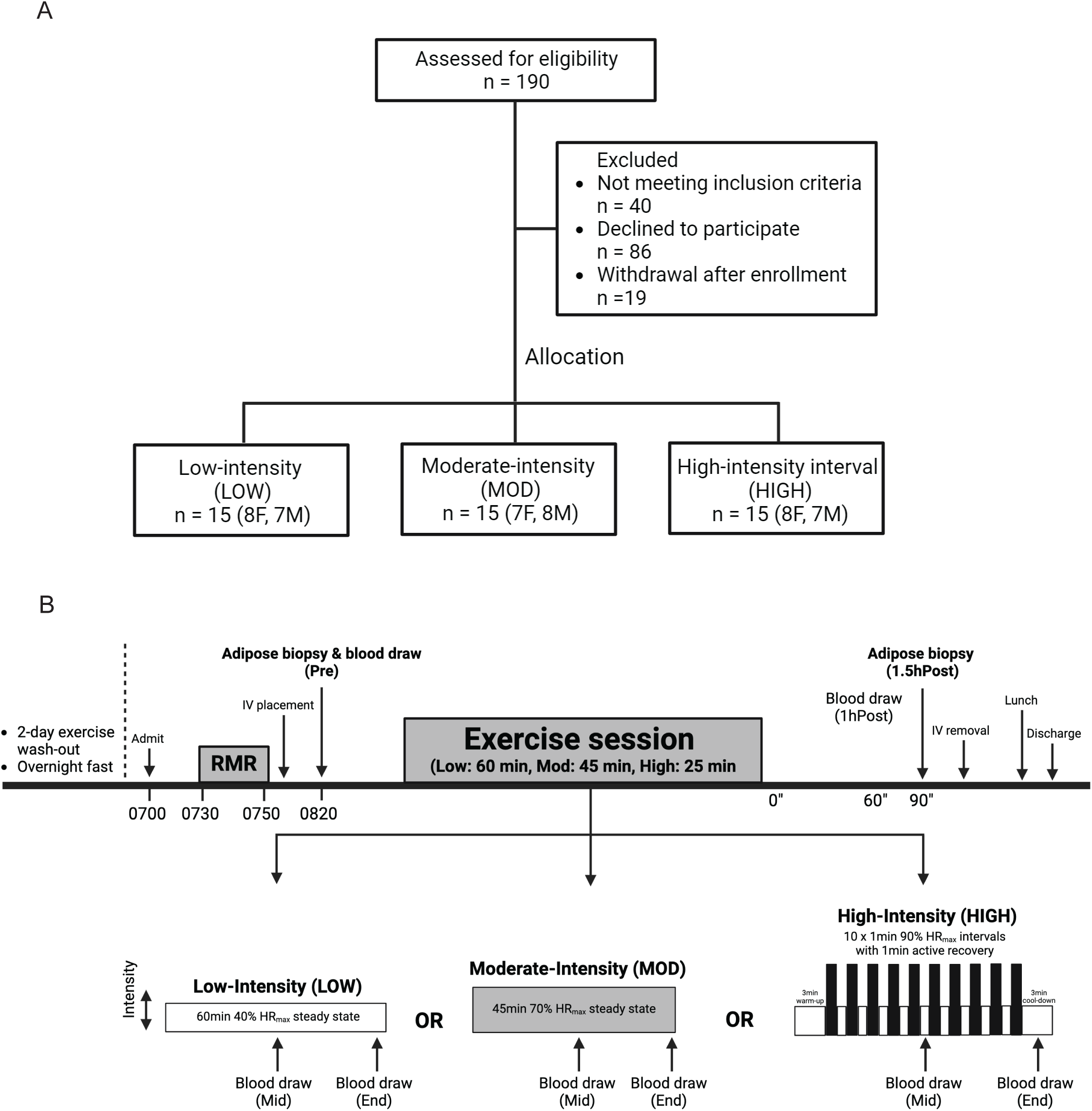
CONSORT diagram and study design. A) CONSORT diagram of the study. B) Schematic of study design.

### Group assignment

Subjects were assigned to one of three exercise treatment groups using a counter-balanced approach to ensure similar distributions of age, sex, baseline body weight, BMI, and peak oxygen consumption (VO_2_peak) across groups. The three exercise treatment groups were: 1) Low-intensity exercise (LOW, 60 min of steady-state exercise at ∼30% VO_2_peak, n=15); 2) Moderate-intensity exercise (MOD, 45 min of steady-state exercise at ∼65% VO_2_peak, n=15); High-intensity exercise (HIGH, 10 x 1 min intervals at ∼90% VO_2_peak with 1 min of active recovery between intervals, n=15).

### Preliminary procedures

#### Body composition

Body composition was determined by dual-energy x-ray absorptiometry (Lunar DPX DEXA scanner, GE, WI).

#### Aerobic capacity (VO_2_peak)

Graded exercise testing was performed on a stationary cycle ergometer (Corvival, Lode, Netherlands) using an incremental exercise protocol starting at 40W and increasing 20W per minute until volitional exhaustion. The rate of oxygen consumption was measured throughout this test using a metabolic cart (Quark CPET, COSMED, Italy) and VO_2 peak_ was determined as the highest 30-second average before volitional fatigue. Measurements of respiratory exchange ratio (RER) ≥ 1.2 and maximal heart rate (HR_max_) ≥ 90% of age-predicted values (i.e., 220-age) were used as secondary indices to help confirm maximal effort during these tests. The experimental exercise sessions were performed on the same cycle ergometer as VO_2_peak was measured.

#### Exercise familiarization exercise

All participants underwent at least one supervised familiarization exercise session during which they performed the exercise protocol consistent with for their group assignment. This familiarization exercise session was completed at least four days before the experimental trial.

### Experimental trial

The experimental trial is outlined in Figure 1B. The evening before the trial, subjects ate a standardized dinner (30% of estimated total daily energy expenditure) at ∼1900h and snack (10% of estimated total daily energy expenditure) at ∼2200h. The next morning, subjects arrived at the clinical facility at 0700h after an overnight fast. After 30 min of resting quietly, resting metabolic rate was measured by indirect calorimetry (TrueOne 2400, ParvoMedics, USA). An intravenous catheter was then inserted into the antecubital vein on one arm for blood collection. At ∼0900 h, an aSAT sample was collected by aspiration 5-10cm distal to the umbilicus, as described previously (16). aSAT samples were snap-frozen in liquid nitrogen and stored at −80°C for later quantification of transcriptome (RNA sequencing) and protein abundance (targeted immunoblots) (see details below). A pre-exercise blood sample was collected in conjunction with the aSAT biopsy. Subjects then performed their prescribed exercise session.

Subjects assigned to LOW performed steady-state exercise at 40% heart rate reserve (HRR) (∼30% VO_2_peak) (17) for 60 min (Figure 1B), subjects assigned to MOD performed steady-state exercise at 70% HRR (∼65% VO_2_peak) for 45 min (Figure 1B), and subjects assigned to HIGH performed 10 x 1 min intervals at 90% HRR (∼90% VO_2_peak) with 1 min of active recovery between intervals (Figure 1B). The HIGH exercise session also included a 3 min warm-up and cool-down before and after the interval protocol (total exercise time in HIGH was 25 min). Exercise intensity was monitored by a telemetry heart rate device (Polar, Finland).

Blood samples were collected at the mid-point (MID) of each exercise session (at ∼30” in LOW, at ∼22” in MOD, and between 5^th^ and 6^th^ interval for HIGH). Additional blood samples were collected during the final 30-60 seconds of exercise (End) and 1hour post-exercise (1hPost).

Blood samples were centrifuged at 2,000 g at 4°C for 15 min. Serum and plasma were aliquoted and stored at −80°C until analysis for circulating factors. After exercise, all subjects rested quietly on a bed for 1h 30min. The post-exercise aSAT biopsy samples (Post) were collected in the same manner as the pre-exercise samples but on the opposite side of the umbilicus.

### Analytical procedures

#### RNA-sequencing

RNA was extracted by using a Qiagen Mini RNA extraction kit (74004, Qiagen). Total RNAs were polyA enriched, and directional RNA-seq libraries were prepared using the NEBNext Ultra II RNA library prep Kit. Paired-end sequencing was conducted at a NovaSeq 6000 sequencer at the Oklahoma Medical Research Foundation (OMRF) Clinical Genomics Center. The paired-end RNA-seq reads were mapped to human transcripts annotated in GENCODE v44 (18) using Bowtie (19). Gene level read counts were estimated using rSeq (20, 21).

#### Bioinformatics

##### Differential analysis

Normalization of gene count data and differential analysis was performed by using DESeq2 (22). Multivariate adaptive shrinkage (mash (23)), an empirical Bayes hierarchical model, was then applied to refine effect estimates across exercise types. Endpoints from mash: posterior effect estimates and local false sign rate (lfsr) represent log fold change and false discovery rate (FDR). lsfr < 0.05 was used as a cutoff for the determination of differentially expressed genes (DEG).

##### Gene set testing

Overrepresentation analysis (ORA) was conducted on DEGs using hypeR R package. Reference gene sets were combined from Gene Ontology, REACTOME, WikiPathways, and Kyoto Encyclopedia of Genes and Genomes (KEGG). Competitive gene set test accounting for inter-gene correlation (CAMERA) (24) was used on all genes, using signed −log(p-value) derived from DESeq2. Reference gene sets were combined from HALLMARK, KEGG and Gene Ontology. BH-adjusted p-value < 0.05 was used as a cutoff for the determination of significantly enriched gene sets. TMSig R package was used for heatmap visualization.

##### Weighted Gene Co-expression Network Analysis (WGCNA)

WGCNA (version 1.73) (25) was employed to identify distinct, non-overlapping clusters (modules) of ASAT transcripts at baseline (n=45 samples). Connectivity for each transcript was calculated by summing its correlation strengths with all other transcripts. A scale-free signed network was constructed using an optimal soft-threshold power (β=6). Using a dynamic tree-cutting algorithm and a merging threshold of 0.3, 16 distinct modules were identified, including Module 0. Module 0, often referred to as the ‘grey’ module, comprises transcripts that could not be assigned to any other module due to weak correlation patterns. Top hub genes within each module were determined based on their kME values (module eigengene-based connectivity). Module eigengenes (MEs), representing the first principal eigenvector of each module, were extracted, and their relationships with clinical traits and secreted protein abundances were evaluated using biweight midcorrelation (26). The coefficient, bicor, is similar to Pearson’s correlation coefficient r, but it produces more robust outcomes by accounting for outliers. Post-exercise ME was acquired by recalculating the first principal component by using the same member genes identified from the pre-exercise modules. Biweight midcorrelation was used to correlate the change of ME (Δ, from pre-exercise to 1.5hPost-exercise) and change of circulating factor concentrations (Δ, from pre-exercise to end of exercise).

##### Cell-type enrichment and deconvolution

Cell-type marker gene lists were curated from three previously published single-nuclei (sn) RNAseq datasets of human adipose tissue (27–29). For gene markers obtained from Emont et al. and Whytock et al., we applied a logFC > 1 threshold to retain genes that were significantly and specifically expressed by each cell type. No expression-level filter was applied to markers from Backdahl et al., as gene expression from spatial transcriptomics is inherently low. The curated markers for each cell type were treated as independent gene sets. Statistical gene set testing was performed using CAMERA. Deconvolution analysis was conducted using a specialized pipeline designed to estimate cell-type proportions (%) in aSAT (30). This pipeline utilizes a gene signature matrix derived from the snRNAseq dataset reported by Whytock et al. (29), which achieved the highest gene coverage per nucleus for aSAT to date, enabled by full-length transcriptomics. Only snRNAseq data from younger adults (18-40 years, n=10) were used to closely match the age range of subjects in this study.

#### Targeted immunoblots

We used capillary electrophoresis-based Western blot (JESS, ProteinSimple, San Jose, CA) to measure the abundance of proteins of interest in aSAT lysates. A portion of each aspirated adipose biopsy sample (∼90 mg wet weight) was homogenized in ice-cold 1X RIPA buffer (89901, Thermofisher) with freshly added protease and phosphatase inhibitors (P8340, P5726, and P0044; Sigma) using two 5 mm steel beads (TissueLyser II, Qiagen, CA). Homogenates were rotated at 50 rpm for 60 min at 4°C and then centrifuged at 4°C for 3 x 15 minutes at 15,000g. Protein concentration was assessed using the bicinchoninic acid method (#23225, Thermofisher) after removing the supernatant. Samples were prepared in 4x Laemmli buffer, heated for 7 minutes at 95°C. Equal amounts of protein (0.16µg) were mixed with the Simple Western sample buffer and fluorescent mix and loaded in capillaries in 12-230 kDa JESS separation module, 25 capillary cartridges (SW-W003). All experiments were performed on the automated JESS device in accordance with the manufacturer’s instructions. Primary antibodies used were Hormone-sensitive lipase (HSL, #18381, Cell Signaling Technology), phospho-HSL (Ser565) (pHSL^S565^, #4137, Cell Signaling Technology), phospho-HSL (Ser660) (pHSL^S660^, #4126, Cell Signaling Technology), Protein kinase B (AKT, #9272, Cell Signaling Technology), phospho-AKT (Ser473) (pAKT^S473^, #9271, Cell Signaling Technology), phospho-AKT (Thr308) (pAKT^T308^, #13038, Cell Signaling Technology), p38 mitogen-activated protein kinase (P38, #9212, Cell Signaling Technology), phospho-P38 MAPK (Thr180/Tyr182) (pP38^T180/Y192^, #9211, Cell Signaling Technology), p44/42 MAPK extracellular signal-regulated kinase (ERK, #4695, Cell Signaling Technology), phospho-p44/42 MAPK (Thr202/Tyr204) (pERK^T202/Y204^, #4376, Cell Signaling Technology), Signal transducer and activator of transcription 3 (STAT3, #12640, Cell Signaling Technology), and phospho-STAT3 (Tyr705) (pSTAT3^Y705^, #9145, Cell Signaling Technology).

#### Blood measurements

Plasma concentrations of glucose (TR-15221, ThermoFisher), plasma fatty acids (NC9517309, NC9517311; Fujifilm Medical Systems), and plasma glycerol (F6428, Sigma) were assessed using commercially available kits. Plasma lactate was measured as previously described (31). Serum insulin concentration was assessed using a chemiluminescent immunoassay (Siemens 1000). Epinephrine and norepinephrine (Abnova, Taipei City,Taiwan; KA1877), and cortisol (R&D Cat. No. KGE008B) were measured by ELISA. Plasma concentrations of IL1β, IL10, IL6, IFNγ, and TNFα were measured using customized Luminex Multiplex kit (HSTCMAG-28SK).

#### Plasma volume correction

A single session of aerobic exercise induces changes in the concentration of many circulating factors (i.e., metabolites or hormones) (32). However, the true effect of exercise on inducing the bona fide synthesis of these factors can be confounded when blood volume is not accounted for, because hemoconcentration during exercise leads to the reduction of plasma volume (33). Therefore, we measured plasma calcium (Ca) concentration as the marker for hemoconcentration (34). The concentration of circulating parameters at Mid, End, 1hPost, and 1.5hPost was corrected for plasma volume as follows: [parameter]_c_ = [parameter]_u_ / (1 + ΔCa(%)/100), where ΔCa(%) refers to the percentage change of Ca concentration relative to pre-exercise (Pre) level and c and u sub-indices denote corrected and uncorrected concentration, respectively.

### Statistical analyses

Two-way ANOVA linear mixed model was used to compare results from all variables except RNA-sequencing data. Post-hoc Tukey test was applied when significant interaction effect was detected. Statistical analysis was done by using R. P-value < 0.05 was considered statistically significant. Data are presented as mean ± SD unless noted otherwise. All statistical analyses were conducted with R (R, Vienna, Austria).

## RESULTS

### Subject characteristics and heart rate responses during exercise

A total of 45 subjects (15 LOW, 15 MOD, and 15 HIGH) completed the study. All subjects were healthy, without obesity, and all were regular exercisers (Table 1). As designed, there was no difference in baseline anthropometric characteristics or aerobic fitness (VO_2_peak) among groups (Table 1). Subjects exhibited a normal range of metabolic health indices, as evidenced by relatively low fasting plasma insulin (5.8 ± 2.9 mU/L), glucose (4.9 ± 0.5 mmol/L), fatty acid (307 ± 151 µmol/L), and triglyceride concentrations (0.8 ± 0.4 mmol/L), all of which did not differ among groups (Table 1). During the steady-state exercise sessions, average HR was 105±7 bpm during LOW and 140±9 bpm during MOD (representing 40±5% and 69±6% HRR, respectively). During HIGH, average HR during the one-minute high-intensity intervals was 161±13 bpm (86±8% HRR and 90±5% HR_max_).

**Table 1.** Baseline subject characteristics and circulating metabolic biomarkers. BMI, Body mass index; SBP, Systolic blood pressure; DBP, Diastolic blood pressure; VO_2,_ Volume of oxygen consumption; RMR, Resting metabolic rate.

|  | **LOW**<br>**(n=15)** | **MOD**<br>**(n=15)** | **HIGH**<br>**(n=15)** | Difference<br>among<br>groups |
| --- | --- | --- | --- | --- |
| Age (years) | 31 ± 6 | 31 ± 6 | 30 ± 7 | NS |
| Sex | 8F, 7M | 7F, 8M | 8F, 7M | NS |
| Height (m) | 1.72 ± 8.0 | 1.75 ± 8.8 | 1.73 ± 9.3 | NS |
| Body mass (kg) | 72.8 ± 9.0 | 74.5 ± 10.7 | 74.0 ± 11.2 | NS |
| Body fat (%) | 26 ± 6 | 26 ± 8 | 26 ± 8 | NS |
| Fat mass (kg) | 17.3 ± 6.1 | 17.3 ± 7.4 | 18.6 ± 7.3 | NS |
| Lean mass (LM) (kg) | 51.5 ± 8.2 | 52.9 ± 10.6 | 52.7 ± 10.8 | NS |
| BMI (kg/m<sup>2</sup>) | 24.6 ± 2.2 | 24.4 ± 2.3 | 24.8 ± 2.6 | NS |
| SBP (mmHg) | 118 ± 8 | 124 ± 9 | 120 ± 11 | NS |
| DBP (mmHg) | 72 ± 7 | 78 ± 8 | 72 ± 9 | NS |
| VO<sub>2</sub> peak (L/min) | 2.6 ± 0.4 | 2.8 ± 0.6 | 2.9 ± 0.6 | NS |
| VO<sub>2</sub> peak (mL/kg/min) | 35.9 ± 4.5 | 37.4 ± 6.3 | 39.0 ± 5.6 | NS |
| VO<sub>2</sub> peak (mL/kg LM/min) | 50.8 ± 4.9 | 52.8 ± 6.5 | 55.1 ± 6.5 | NS |
| Fat oxidation (μmol/min) | 352 ± 79 | 349 ± 92 | 363 ± 73 | NS |
| RMR (kcal/day) | 1432 ± 160 | 1460 ± 250 | 1526 ± 244 | NS |
| Fasting insulin (mU/L) | 6.3 ± 2.7 | 5.4 ± 2.5 | 5.6 ± 3.5 | NS |
| Fasting glucose (mmol/L) | 4.9 ± 0.7 | 5.1 ± 0.5 | 4.86 ± 0.5 | NS |
| Fasting fatty acids (μM) | 361 ± 200 | 283 ± 125 | 278 ± 106 | NS |
| Fasting triglycerides (mM) | 0.77 ± 0.28 | 0.89 ± 0.41 | 0.84 ± 0.40 | NS |

### Concentrations of circulating factors

Plasma concentrations of all circulating factors measured in our study were corrected for the rapid reduction in plasma volume that occurs at the onset of exercise, using calcium as a marker of this exercise-induced hemoconcentration (34) (Figure S1A). As anticipated, exercise significantly increased plasma epinephrine and norepinephrine concentrations above Pre in all groups (all p<0.001), and concentrations of both hormones were more than 2-fold greater in HIGH compared with LOW and MOD at the end of exercise (p<0.001), with no difference between LOW and MOD (Figure 2A, 2B). Similarly, exercise significantly increased plasma lactate concentrations above Pre in all groups (all P<0.001), with the highest concentrations found after HIGH (P<0.001) vs. MOD and LOW; Figure 2C). Plasma lactate concentration during MOD was also significantly greater than LOW (P<0.01) (Figure 2C). Plasma concentrations of epinephrine and lactate returned to pre-exercise levels 1h after exercise (1hPost) in all groups. Plasma norepinephrine concentration also declined to near pre-exercise level at 1hPost in all groups, however, 1hPost values remained slightly, yet significantly higher than Pre in both LOW and MOD (both p≤0.05), but not in HIGH (Figure 2B). In contrast to plasma catecholamine and lactate concentrations, we did not detect a significant change in plasma cortisol concentration during exercise in any of the groups (Figure 2D).

**Figure 2.**
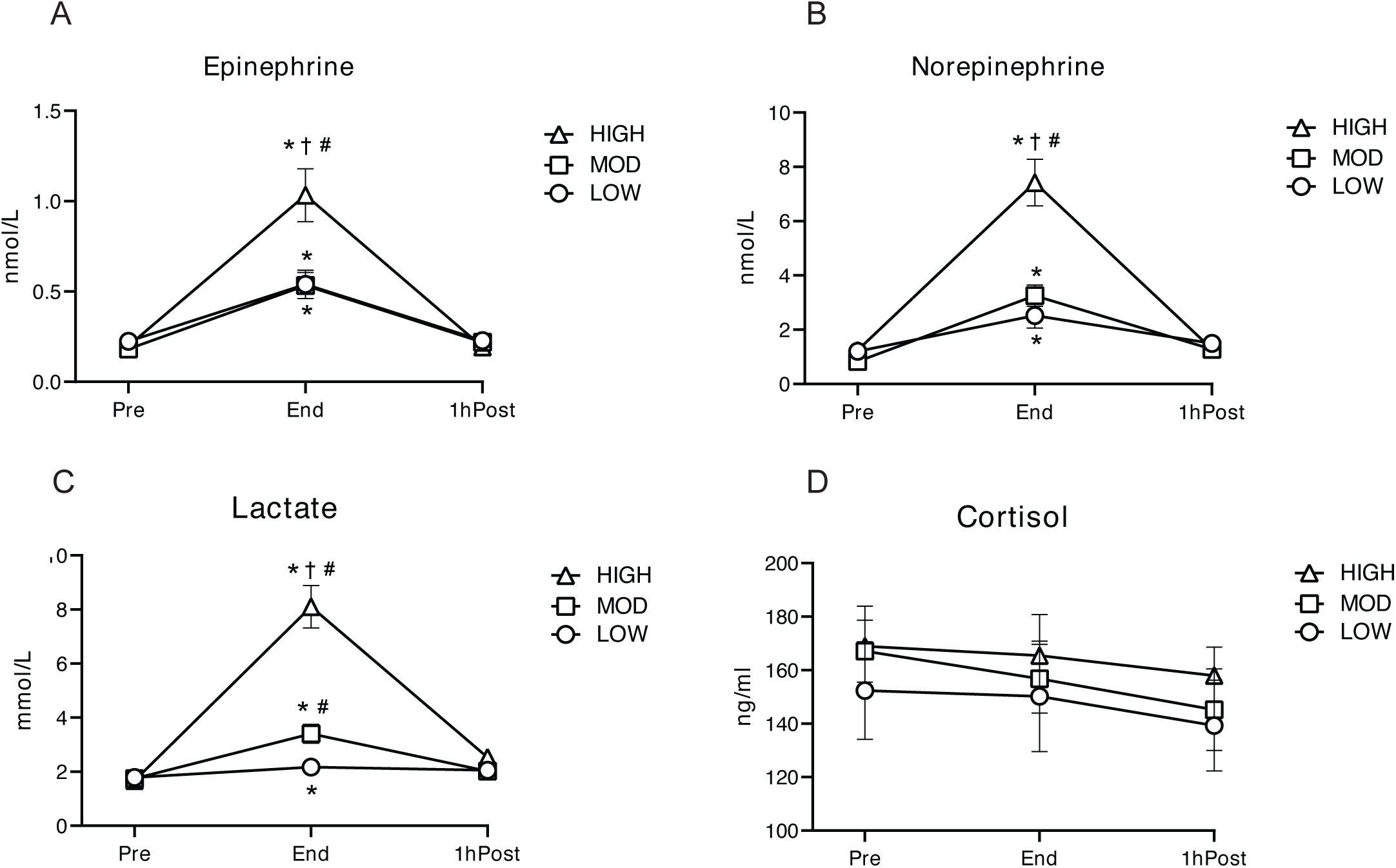
Concentrations of circulating factors during and post LOW, MOD, and HIGH. A) Plasma epinephrine concentration. B) Plasma norepinephrine concentration. C) Plasma lactate concentration. D) Plasma cortisol concentration. *p<0.05 vs. Pre; †p<0.05 vs. MOD; #p<0.05 vs. LOW. Significant overall Time x Group interaction effects were detected in epinephrine, norepinephrine, and lactate. (p<0.001, Type III ANOVA). Detailed p-values are included in the Results. Data is presented as Mean±SEM.

As expected, plasma glycerol concentration increased during exercise compared with Pre in all groups (main effect of time p<0.0001 for both Mid and End), and concentrations declined to near pre-exercise levels 1hPost exercise (Figure S1B). There were no significant differences in plasma glycerol concentrations among groups (Figure S1B). Plasma fatty acid concentration was significantly elevated after exercise in LOW (p<0.0001 and p<0.002 for End and 1hPost, respectively) (Figure S1B). During MOD, plasma fatty acid concentration was not significantly elevated above pre-exercise and remained less than LOW after exercise (p=0.001 and p=0.036 for End and 1hPost, respectively) (Figure S1B). In contrast to LOW and MOD, HIGH reduced plasma fatty acid concentration during exercise (p<0.05), and fatty acid concentration 1hPost in HIGH was significantly lower than both MOD (p=0.013) and LOW (p<0.0001) (Figure S1B). Plasma insulin concentration was slightly, yet significantly elevated at 1hPost in HIGH (p<0.05), while glucose concentrations were unaltered by all exercise groups (Figure S1C). Exercise induced a modest increase in plasma triglyceride concentration (main effect of time; p<0.05 for both Mid and End), and returned to the pre-exercise level at 1hPost (Figure S1C).

When accounting for the exercise-induced reduction in plasma volume, exercise did not significantly alter plasma concentrations of IL1β, IL6, IFNγ, or TNFα (Figure S2A), but there was a main effect of exercise on the increase in IL10 (p<0.05) (Figure S2E). In contrast, without adjusting for plasma volume, we found a significant main effect of exercise on the concentrations of IL1β, IFNγ, TNFα, and IL10 at End (all p<0.05) (Figure S2B) except for IL6. Collectively, these findings suggest that the increase in circulating cytokine bioavailability may be largely attributed to exercise-induced hemoconcentration.

### Acute LOW, MOD, and HIGH induced distinct transcriptomic responses in aSAT

A total of 17,466 genes were detected by our RNAseq analysis and comparing aSAT samples collected before exercise (Pre) and 1.5h after exercise (Post). We applied multivariate adaptive shrinkage (mash (23)), an empirical Bayes hierarchical model, following differential analysis with DESeq2 to refine effect estimates across exercise types. Mash provides posterior effect estimates and the local false sign rate (lfsr), a stricter alternative to FDR that ensures effects are both significant and correctly signed. We found 2671 differentially expressed genes (DEG) after LOW (1494 upregulated and 1177 downregulated), 2974 DEGs after MOD (1245 upregulated and 1729 downregulated), and 2570 DEGs after HIGH (1379 upregulated and 1191 downregulated) (lsfr<0.05) (Figure 3A). Overrepresentation analysis (ORA) suggested that inflammatory responses, MAPK cascade, and vasculature development were globally enriched from DEGs across groups (adjusted p<0.05, Figure 3B). Notably, most of enriched pathways by HIGH were driven by upregulated DEGs, which included immune cell activation, cytoskeleton organization, circadian rhythm, and insulin signaling (adjusted p<0.05, Figure 3B). Additionally, competitive geneset tests (CAMERA) (24) using all genes also identified distinct sets of biological pathways significantly altered by each of the three exercise treatments. For example, the top-upregulated pathways after LOW included protein translation and immune activation pathways, while pathways related to adipogenesis and lipid metabolism were downregulated (adjusted p<0.05, Figure 3SA). The upregulated pathways in MOD included protein translation, ribosome biogenesis, and mitochondrial metabolism, while ECM protein organization and β-catenin binding were downregulated (adjusted p<0.05, Figure S3A). Interestingly, many DEGs were group-specific and exhibited opposing regulation across exercise intensities. For example, 394 DEGs downregulated in LOW and MOD were upregulated in HIGH, with overrepresentation in cellular signaling, regulation in gene expression, cytokine production, and response to peptide (adjusted p<0.05, Figure 3C), suggesting distinct transcriptional effect by HIGH. This was supported by principal component analysis (PCA), which revealed distinct gene expression patterns (Δ1.5hPost-Pre) at the sample level, particularly in the HIGH compared with MOD and LOW (Figure S3B). Conversely, we observed that DEGs (n=324) enriched for MAPK cascade and protein metabolism were upregulated by HIGH and LOW, but downregulated by MOD (adjusted p<0.05, Figure 3C).

**Figure 3.**
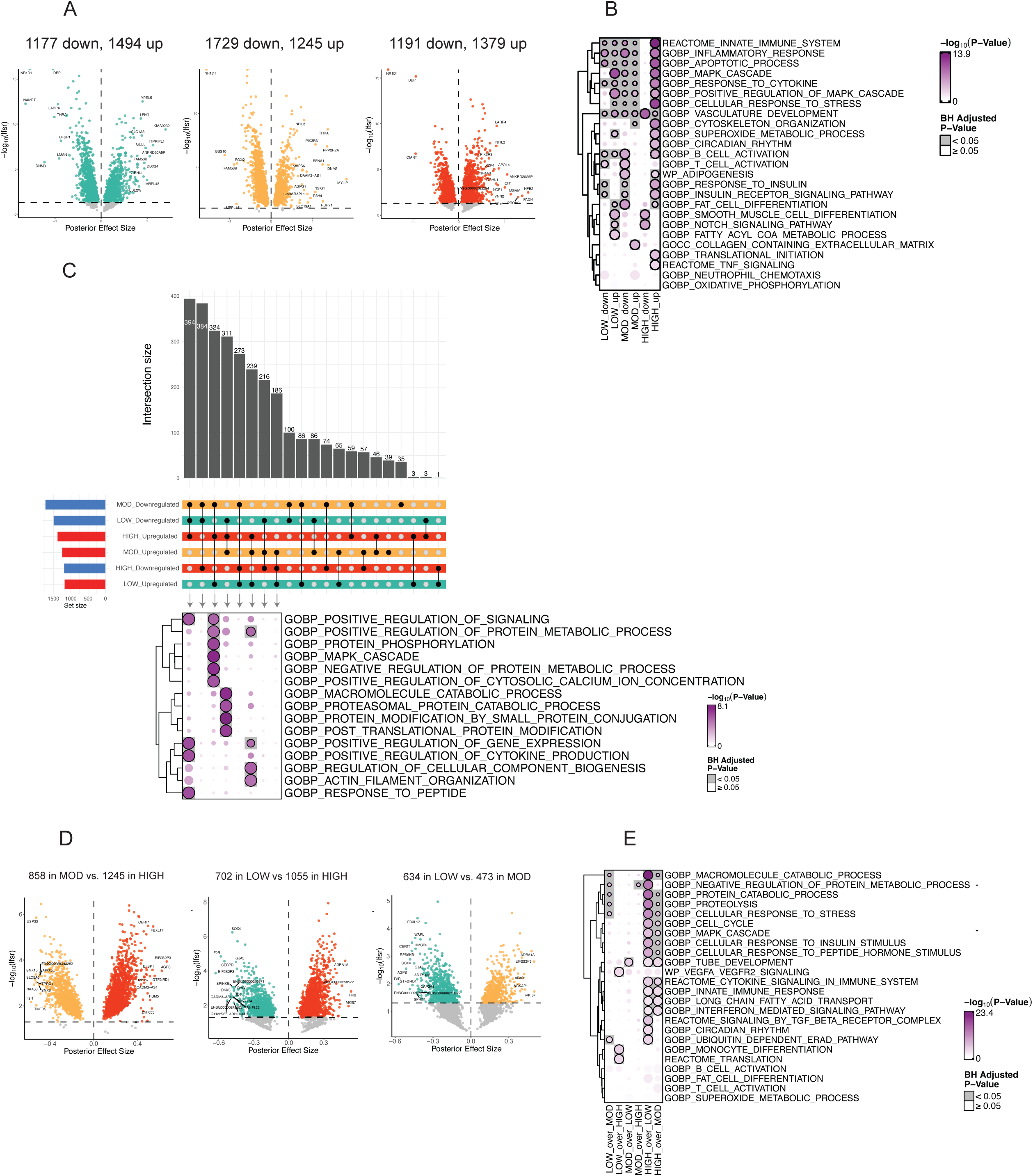
Transcriptomic responses in aSAT 1.5hour after LOW, MOD, and HIGH. A) Volcano plots of DEGs in LOW, MOD, and HIGH 1hPost vs. Pre (lfsr<0.05). B) ORA results from upregulated and downregulated DEGs for each exercise group. C) Upset plots of upregulated and downregulated DEGs after LOW, MOD, and HIGH. Black dots indicate the corresponding bar plots of DEG count. Intersecting lines indicate shared DEGs among exercise groups. Blue bar represents downregulation and red bar represents upregulation. ORA was conducted on intersecting DEGs across three exercise groups. D) Volcano plots of DEGs comparing the acute effects of HIGH vs. MOD, HIGH vs. LOW, and MOD vs. LOW (lsfr<0.05). Colored DEGs represents larger effect by corresponding exercise treatment. E) ORA was performed on DEGs identified from comparisons between different acute exercise treatments. DEG, Differentially expressed genes; lfsr, local false sign rate; ORA, Overrepresentation analysis; GOBP, Gene Ontology-Biological Process; WP; WikiPathways

To further investigate the effects of exercise type on the aSAT transcriptome, we compared DEGs across our three different exercise treatments. We identified 2103 DEGs in MOD vs. HIGH (1245 greater in HIGH, 858 greater in MOD), 1757 DEGs in LOW vs. HIGH (1055 greater in HIGH, 702 greater in LOW), and 1107 DEGs in LOW vs. MOD (473 greater in MOD, 634 greater in LOW) (all lfsr < 0.05) (Figure 3D). Notably, ORA revealed that pathways involved in protein catabolism, cell cycle, response to external stimulus, and inflammation were enriched by HIGH compared with MOD and LOW (adjusted p<0.05, Figure 3E), indicating that these biological pathways might require higher exercise intensity to occur. CAMERA additionally suggested ribosome biogenesis and oxidative phosphorylation were more pronounced in MOD compared with HIGH while cytoskeleton remodeling and calcium channel activity were more strongly induced by LOW compared with MOD (adjusted p<0.05, Figure S3C).

### Circadian clock genes are strongly altered by acute exercise

Although the shared downregulated DEGs among three exercise groups was the 2^nd^ largest intersection (n=384, Figure 3C), ORA was unable to detect enriched pathways. Therefore, we manually inspected these DEGs to examine genes that were suppressed by all three exercise groups. We found that many circadian clock genes were strongly downregulated across all treatments. Notably, a core clock gene Nuclear Receptor Subfamily 1 Group D Member 1 (*NR1D1*; also referred to as Rev-Erbα) that inhibits the transcription of master clock gene *BMAL1*, was the most significantly downregulated gene in all exercise groups (Figure 4A). Other core clock genes such as Circadian-Associated Repressor of Transcription (*CIART*), D-box binding PAR bZIP transcription factor (*DBP*), and Period Circadian Regulator 1 (*PER1*) were also suppressed by exercise. Conversely, Nicotinamide Phosphoribosyltransferase (*NAMPT*), also known to regulate circadian rhythm (35) was upregulated by all treatments in most subjects (Figure 4A). We did not observe any exercise-induced changes in the gene expression of *BMAL1* or *CLOCK* (Figure 4B), which are considered the ‘master clock’ transcription factors that regulate the expression of other circadian genes (36).

**Figure 4.**
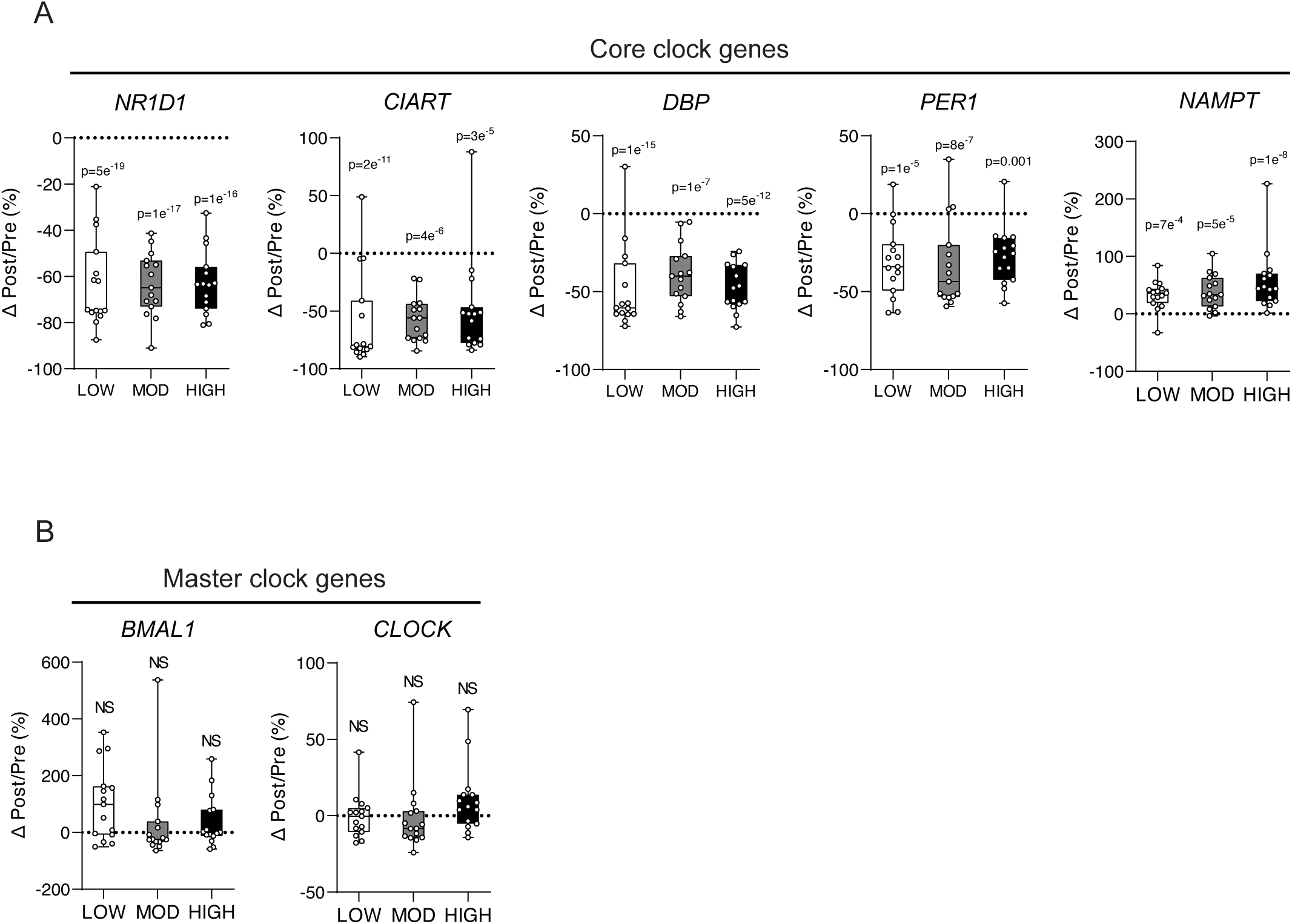
Alterations in circadian clock genes in aSAT 1.5hour after exercise. A) Percentage change of core clock genes (Δ1hPost/Pre) after LOW, MOD, and HIGH. B) Percentage change of master clock genes (Δ1hPost/Pre) after LOW, MOD, and HIGH. P-values represent the BH-corrected adjusted p-value obtained from DESeq2 analysis. NS, Not significant.

### Gene modules identified by WGCNA are linked to clinical traits

We conducted weighted gene co-expression network analysis (WGCNA) on pre-exercise transcripts to identify gene clusters (i.e., modules) and their module eigengene (ME; first principal component of module that represents overall gene expression of the module) that may help identify how genes are co-expressed and how they may be functionally related. A total of 16 modules were identified (including the ‘grey’ module, Module 0), each containing at least 200 genes. ORA revealed distinct biological pathways associated with each module (Figure 5A). For example, ME1 was enriched in pathways related to aerobic respiration, small molecule metabolic process, and lipid metabolism (adjusted p<0.05, Figures 5A), and its ‘hub genes’ (i.e., highly connected genes within ME) included mitochondrial genes (e.g., *SDHB, NDUF, COX*) (Figure S4A). Next, we correlated MEs with clinical, systemic, and histological traits (i.e., adipocyte size, collagen abundance (assessed by Sirius Red), CD14^+^ and CD206^+^ macrophages; Figure 5B, S3B), and we also examined the module membership of DEGs (Figure 5C) to assess the potential link between these gene modules and their biological function, as well as the effects of acute exercise. For example, ME1 was linked to females, inversely associated with adiposity and fat cell size (FCS) and ∼13% of member genes were differentially regulated by exercise (Figure 5B, 5C). ME8 was enriched for inflammation, cell migration, and lipid metabolism (adjusted p<0.05, Figure 5A), with its top 10 hub genes being key factors in immune cell activation, migration, and cytokine production (Figure S4A). Additionally, ME8 was positively associated with adiposity, waist circumference, fasting glucose concentration, adipocyte size (p<0.05, Figure 5B), and HOMA-IR, a marker of whole-body insulin resistance (IR) (p<0.05, Figure 5B), with more than 50% of module member genes being differentially regulated by exercise (Figure 5C).

**Figure 5.**
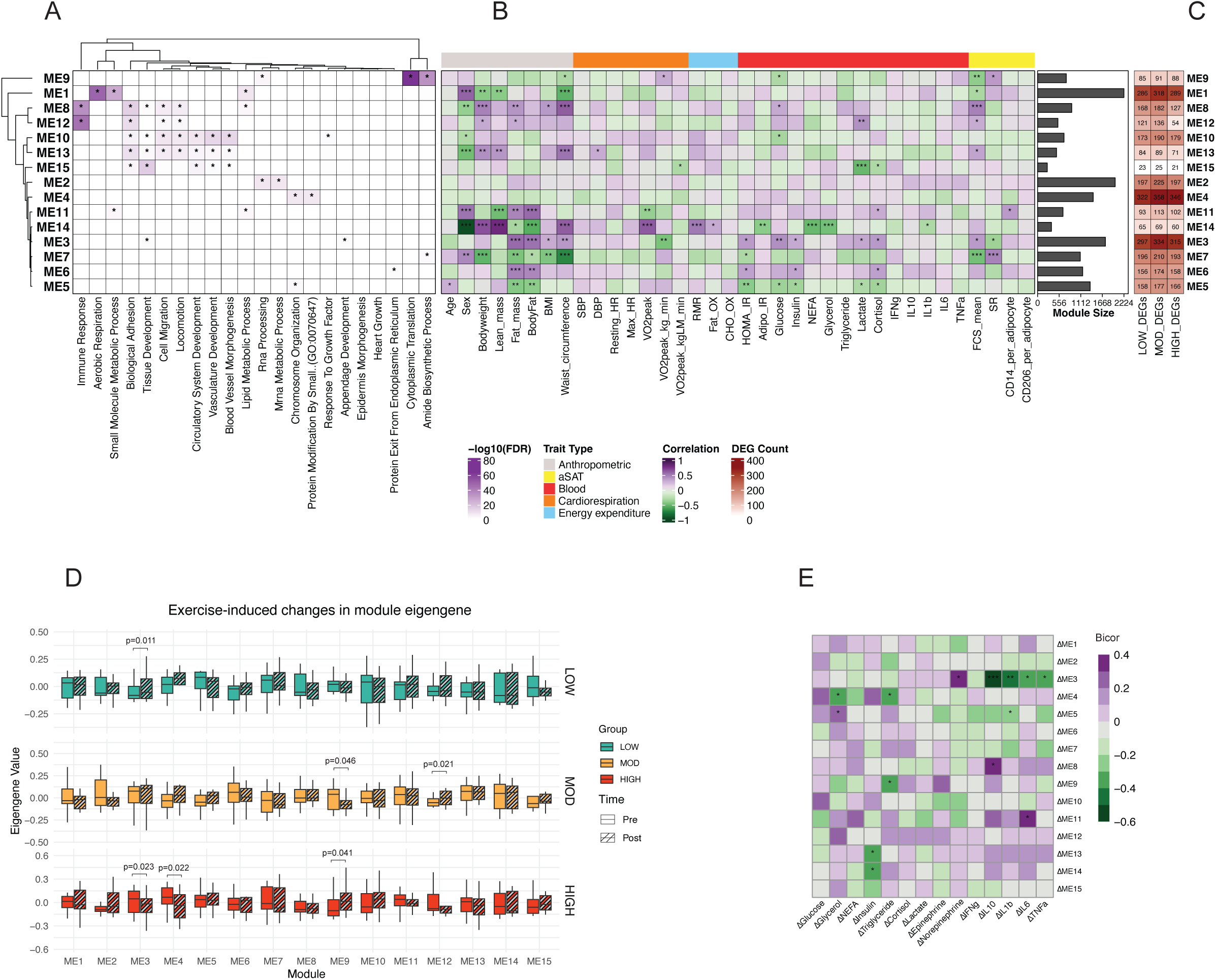
Integration of transcriptomics with clinical/tissue traits. WGCNA was performed on the pre-exercise transcriptomics data (n=45). A) ORA was conducted on module member genes. Significantly overrepresented terms are marked with asterisk (FDR<0.05). B) ME was correlated with clinical and sub-clinical traits using biweight midcorrelation. *p<0.05; **p<0.01; ***p<0.001. Module size (number of genes in each module) is shown in horizontal bar graphs, right to the correlation heatmap. C) DEGs were mapped to module member genes. Count of DEGs is shown in numbers. D) Change of ME after LOW, MOD, and HIGH. F) Correlation of ΔME (1.5hPost/Pre) with Δblood analytes (1hPost/Pre). *p<0.05; **p <0.01; ***p<0.001. Bicor, Biweight midcorrelation r value.

Then, we examined if certain modules may be more responsive to acute exercise. Notably, ME9 was enriched for pathways involved in translation and amide biosynthesis (Figure 5A) and was significantly downregulated by MOD (p=0.048, Figure 5D). Conversely, we found that ME9 was increased in HIGH (p=0.041, Figure 5D). Several other MEs were also impacted by acute exercise: ME3 increased after LOW (p=0.011), ME12 increased after MOD (p=0.021), and ME3 and ME4 decreased after HIGH (p=0.023, 0.022, respectively) (Figure 5D).

To explore potential transcriptional regulation in aSAT by circulating factors altered during exercise, we correlated the change in unadjusted blood analytes from Pre to End (Δ) with changes in ME from Pre to 1.5hPost (ΔME). ΔInsulin, known for its role in inhibiting adipocyte lipolysis, was negatively associated with ΔME13 and ΔME14 (both p<0.05, Figure 5E). Notably, ΔME3 showed a positive correlation with Δnorepinephrine but an inverse relationship with several circulating cytokines (e.g., ΔIL10, ΔIL1β, ΔIL6, ΔTNFα), suggesting that genes in ME3 may be transcriptionally regulated by exercise-induced hormonal changes (p<0.05, Figure 5E). Among the top 20 hub genes for ME3, *ANXA1, ANXA7, CCPG1, CRYAB, LNPEP*, and *VLDLR*—spanning immune response, exocytosis, cell cycle regulation, and lipid metabolism— were differentially regulated across exercise groups (lfsr<0.05, Figure S4C). Interestingly, ANXA7, CCPG1, and LNPEP exhibited significant interaction effects among exercise groups (lfsr<0.05, Figure S4D).

### Acute exercise downregulated ERK phosphorylation at 1.5hPost

Targeted analysis revealed that exercise decreased the phosphorylation of ERK in aSAT (expressed as the ratio of protein abundance of pERK^T202/Y204^:total ERK) after exercise in all groups (p<0.05) with no differences observed among groups (Figure 6A). In contrast, we did not detect an effect of exercise on the phosphorylation of P38 (ratio pP38^T180/Y182^:total P38) or STAT3 (ratio pSTAT3^Y705^:total STAT3) in any of the groups (Figure 6B, C). Phosphorylation of one of the chief lipolytic enzymes, HSL on either the AMPK regulatory site serine 565 (expressed as pHSL^S565^:total HSL) or the PKA regulatory site, serine 660 (pHSL^S660^:HSL) were not increased in aSAT collected 1.5 hours after any of the exercise sessions (Figure 6D). This aligns with the plasma epinephrine concentration, which returned to pre-exercise levels 1 hour after exercise in all groups (Figures 2A). Similarly, there were no differences in the phosphorylation of AKT (pAKT^T308^:AKT or pAKT^S473^:AKT) in the aSAT samples collected after LOW, MOD, or HIGH (Figure 6E).

**Figure 6.**
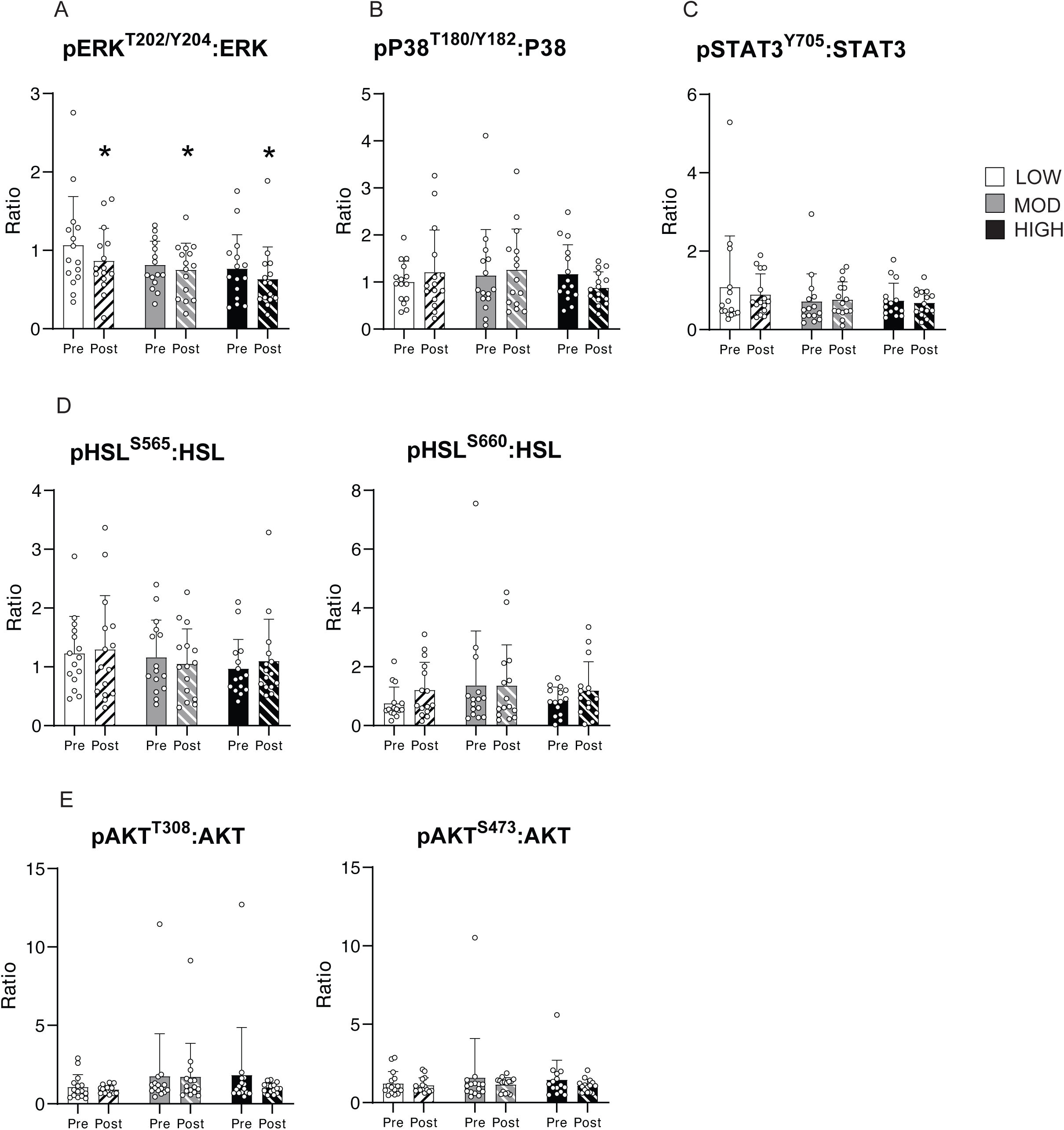
Protein phosphorylation of metabolic proteins in aSAT 1.5hour after LOW, MOD, and HIGH. A) Ratio of pERK^T202/Y204^:ERK. B) Ratio of pP38^T180/Y182^:P38. C) Ratio of pSTAT3^Y705^:STAT3. D) Ratio of pHSL^S565^:HSL and pHSL^S660^:HSL. E) Ratio of pAKT^T308^:AKT and pAKT^S473^:AKT. *p<0.05 vs. Pre. There was no significant Time x Group interaction effect. Data is presented as Mean±SD.

### Integration with single-cell data suggests cell-type dependent responses in aSAT by acute exercise

Using cell-type markers acquired from three previously published reports (27–29), we found a strikingly distinct pattern of transcriptional activation among cell types that were differentially regulated by the three different exercise treatments (Figure 7A). Notably, marker genes for infiltrating mesenchymal immune cells (i.e., macrophage, monocyte, neutrophil) were upregulated in HIGH from all three cell-type marker gene sets (Figure 7A), suggesting that these cells may underlie the strong upregulation of inflammatory pathways observed in HIGH (Figure 3B). Conversely, marker genes for adipocytes were downregulated in LOW (Figure 7A), consistent with the acute downregulation of ‘Adipogenesis’ pathway we found in our RNAseq analysis (Figure S3A). Additionally, we observed a potential exercise intensity-dependent effect on vascular cells, with LOW upregulating gene expression in endothelial and vascular cells, while HIGH exerted the opposite effect (Figure 7A).

**Figure 7.**
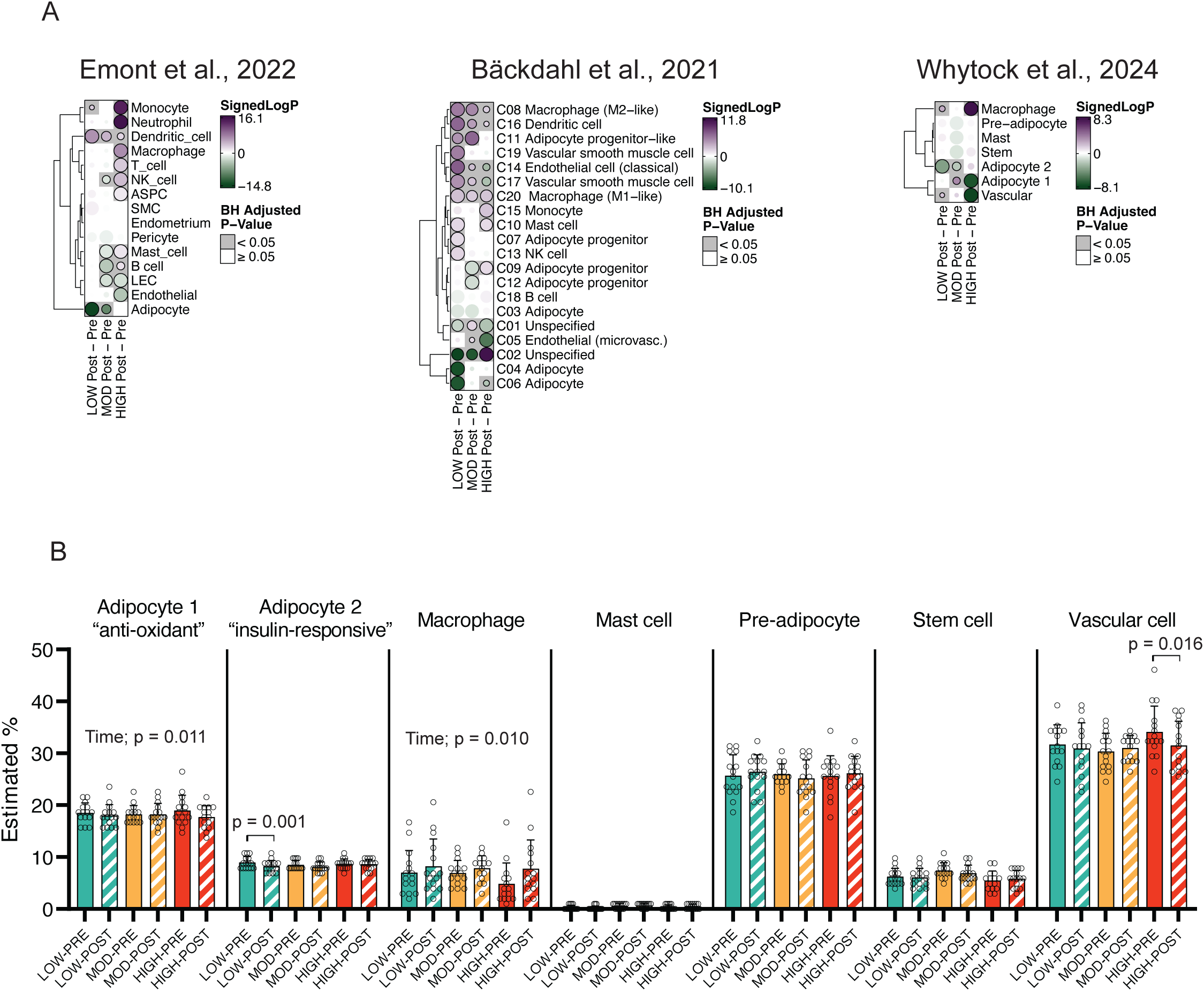
Cell-type specific responses and estimated cell-type proportion of aSAT 1.5hour after LOW, MOD, and HIGH. A) Cell-type enrichment analysis using cell-type markers derived from three publicly available single nuclei RNA sequencing datasets. Competitive gene testing was used to assess cell-type enrichment by exercise. The size of each circle represents the absolute signed log_10_(adjusted p-value). Positive signed log_10_(adjusted p-value) represents activation and negative signed log_10_(adjusted p-value) represents suppression of cell-type gene markers. B) Estimated proportion of cell types via deconvolution analysis using signature matrix from Whytock et al., 2024. Time, Main effect of time. ASPC, Adipose stem and progenitor cells; LEC, Lymphatic endothelial cells; NK, Natural killer; SMC, Smooth muscle cells.

Building on the aSAT single nuclei ‘signature matrix’ from 10 individuals (age 18-40) reported from Whytock et al (29) (see details in the Methods), we applied a recently developed deconvolution algorithm optimized for human aSAT (30) to estimate the proportion (%) of each cell type before and after acute exercise. Deconvolution analysis revealed an increased estimated proportion of macrophages in aSAT 1.5hPost exercise (main effect of time; p=0.01), suggesting the possibility of rapid infiltration of immune cells into aSAT after exercise (Figure 7B). In contrast, the estimated proportion of adipocyte 1, a sub-type of adipocytes that is characterized by high expression of genes involved in anti-oxidant activity (*GPX1* & *GPX4*) (29), was reduced 1.5hPost exercise (main effect of time; p=0.011). The estimated proportion of adipocyte 2, enriched with upregulation of insulin signaling pathways (29), was only reduced by LOW (Figure 7B). Additionally, the estimated proportion of vascular cells was reduced only by HIGH (p=0.016) (Figure 7B). Overall, our integration of single-cell data suggests that early transcriptional and cellular responses in aSAT may be differentially regulated after a session of exercise performed at different intensity and energy expenditure.

## DISCUSSION

The tissue and cellular adaptations that occur with chronic exercise training result from the accumulation of signaling events/responses to each session of exercise (37). In this study, we aimed to discover the early transcriptomic responses in aSAT triggered after a session of exercise that may contribute to the maintenance or remodeling of aSAT milieu. Among the most novel findings of this study were some distinct alterations in aSAT transcriptome observed 1.5 hours after the three different exercise sessions (LOW, MOD, HIGH), all of which are commonly prescribed for maintaining fitness and health. HIGH elicited the strongest upregulation of genes involved in inflammation, protein catabolism, and response to insulin and hormones, whereas MOD and LOW predominantly increased genes associated with ribosomal function and oxidative phosphorylation pathways. Alteration in circadian clock genes was largely consistent across three exercise modalities, accompanied by reduced phosphorylation of MAPK protein and altered circulating cytokine abundances. Our comprehensive bioinformatic analyses provide insights into the clinical relevance of exercise-induced alterations in aSAT transcriptomes and cell types. These molecular modifications in aSAT that occur shortly after each session of exercise may contribute to the maintenance of adipose tissue function in those who exercise regularly.

We recently reported that exercise training induced some structural and proteomic adaptations in aSAT that were independent of changes in fat mass, and these adaptations were linked to improved cardiometabolic health profiles (3, 4). These adaptations included increased vasculature, altered ECM collagen profile, and elevated abundances of ribosomal, mitochondrial, and lipogenic proteins in aSAT (3, 4). In the present study, our assessments of changes in transcriptional pathways that occur shortly after a session of exercise - including rapid alterations in pathways regulating vasculature development, ECM remodeling, ribosome biogenesis, and lipid metabolism - provide valuable insights into the acute effects of exercise on aSAT that may underlie long-term adaptations. Notably, many responses to acute exercise were different among our three exercise treatments. For example, cytokine production was downregulated by MOD and LOW but upregulated by HIGH. Conversely, oxidative phosphorylation and ribosome biogenesis were more enriched by MOD compared with LOW or HIGH, which is consistent with the findings from a previous exercise training study (38). Because the estimated energy expended during MOD (∼400 kcal) was greater than both LOW and HIGH (both ∼250 kcal), it is possible that these responses might be sensitive to factors related to exercise energy expenditure.

Expression of inflammatory genes has been reported to increase in adipose tissue after a session of exercise in both preclinical models (39, 40) and in humans (41–43), suggesting a transient induction of aSAT inflammation may be an intrinsic response to the stress of exercise. Factors responsible for this exercise-induced inflammatory response in aSAT are not completely understood, but an accumulation of non-esterified fatty acids may be an important contributor (44, 45). The robust upregulation of genes involved in inflammatory signaling and immune cell activations (e.g., macrophages, B cell, T cell) that we observed in HIGH, may have been at least partly due to the restriction in adipose tissue blood flow known to occur during high-intensity exercise (46, 47), and the resultant “trapping” of non-esterified fatty acids within the tissue (44, 45), reflected in our study by the relatively low plasma fatty acid concentration during HIGH. Similarly, a relatively high rate of lipolysis relative to energy expenditure during low-to-moderate-intensity exercise (48) may result in an intracellular accumulation of non-esterified fatty acids, which in turn may contribute to the upregulation in aSAT inflammation we observed in LOW and MOD. Collectively, these findings suggest that the upregulation of gene expression in immune and inflammatory pathways in aSAT may be typical response after exercise, but the extent and specific immune cell types involved in this response may be dependent on exercise intensity.

Our novel findings also demonstrate that the expression of genes for several core clock components (e.g., *NR1D1, NAMPT, DBP, PER1, CIART*) (49, 50) were significantly altered by acute exercise. It is well documented that lipid mobilization and storage in aSAT are impacted by circadian regulation (51, 52), and the disruption of the circadian rhythm of these processes has been found to be linked to metabolic health complications (53). Interestingly, the gene we found to have the greatest magnitude of change in response to all three exercise treatments was the reduction in gene expression of the core clock component, *NR1D1* (Rev-Erbα), which was recently reported to play a critical role in adipose tissue abnormalities common in obesity (e.g., fibrotic ECM and macrophage infiltration) (54). Hunter et al. demonstrated that compared with wild-type mice on a high fat diet (HFD), mice with a specific deletion of *NR1D1* in white adipose tissue were protected from the large increase in fibrosis and macrophage infiltration in adipose tissue in response to the same HFD, despite an even greater increase in adiposity in the knockout animals (54). The adipose-specific *NR1D1* knockout mice were also protected from developing insulin resistance in response to the HFD (54). Additionally, the increased gene expression of *NAMPT* that we observed after exercise also supports the notion that acute exercise may trigger cues to favorably modify aSAT because *NAMPT*, a key enzyme that mediates NAD+ biosynthesis and circadian lipid metabolism (55), has been found to be important in the healthy expansion of subcutaneous adipose tissue (56). Because a greater *<u>capacity</u>*to store lipid in subcutaneous adipose tissue can reduce excessive ectopic lipid deposition in tissues such as the liver, skeletal muscle, pancreas, etc, thereby improving cardiometabolic health (57, 58), it is intriguing to consider that exercise-induced changes in the expression of *NR1D1* and *NAMPT* in aSAT after each session of exercise may lead to beneficial adaptations by reducing ectopic lipid deposition and the resultant dysfunction in metabolically active tissues.

The reduction in *PER1* we observed after exercise is in contrast with a recent finding that a session of moderate-to-vigorous exercise (80% VO_2_max) increased the mRNA expression of *PER1* in aSAT after 8-weeks of exercise training in adults with overweight/obesity (59). Reasons for this discrepancy are not clear, but this suggests changes in circadian transcripts after acute exercise may depend on factors such as the degree of adiposity, training-, and/or nutritional-status (59, 60). Interestingly, the expression of ‘master clock’ genes (i.e., *BMAL1* and *CLOCK*) whose transcription is suppressed by *NR1D1* and *PER* (61), did not change after acute exercise in either our study or Dreher et al. (59), which may indicate that the main aSAT clock was stable. We acknowledge it is possible we may have not captured an exercise-induced change in expression of the master clock genes in our samples collected 1.5h after the exercise session. In addition, although our finding of exercise-induced alterations in core clock genes is interesting, because circadian rhythm is tightly associated with energy homeostasis in aSAT (62), we acknowledge these responses may be driven by an indirect effect of the energy deficit induced by exercise, rather than a direct effect of the exercise, per se. However, it was recently reported that an energy restriction weight loss program (without exercise) did not increase mRNA expression of *NR1D1* or *PER1* in adipose tissue from adults with obesity (60, 63) suggesting the significant alterations in core clock genes we observed may be at least partially mediated by a direct effect of the exercise stimulus.

Our findings that plasma concentrations of circulating pro- and anti-inflammatory cytokines were elevated at the end of moderate- and high-intensity exercise aligns with previous studies (64–66). Here, we report that low-intensity exercise (40% VO_2_max) was also sufficient to increase circulating concentration of cytokines. Therefore, despite very different exercise stimuli and energy expenditure during LOW, MOD, and HIGH in our study, the effect of exercise on the circulating cytokines we measured were remarkably similar. However, our findings suggest that the increased concentration of circulating cytokines in response to acute exercise may be largely driven by exercise-induced hemoconcentration rather than de novo secretion, except for IL10, which was the only cytokine we measured that remained significantly elevated after correction for the exercise-induced reduction in plasma volume. It is possible an increased secretion of IL10 from aSAT may have contributed to the elevation in circulating IL10, particularly by high-intensity exercise, given our findings from our RNAseq analysis indicating that cytokine production and IL10 biogenesis pathways were upregulated in HIGH.

We used WGCNA to identify highly correlated gene clusters, which enabled us to probe the possible impact of exercise-induced changes in aSAT transcripts on cardiometabolic health. For instance, in aSAT samples collected before exercise, the positive association between fat mass or adipocyte size and ME-8 and ME-12, modules enriched for inflammatory pathways, corroborates the well-established link between aSAT inflammation and adiposity (67). By integrating WGCNA modules and post-exercise aSAT transcriptomics, we aimed to comprehensively characterize exercise-induced transcriptional responses in aSAT. Among the gene modules that were significantly altered by exercise, the change of ME3, a module that was associated with ‘unfavorable’ cardiometabolic traits (e.g., higher adiposity, adipocyte size, and fasting glucose, insulin, and cortisol), was inversely correlated with changes in circulating cytokines such as IL10, IL1β, IL6, and TNFα. Perhaps, the increased concentrations of these cytokines during exercise may have acted to downregulate the expression of ME3 genes. Collectively, this analysis bridges exercise-induced effects in aSAT with cardiometabolic health traits, providing clinical insights into the role of exercise on transcriptional regulation in aSAT.

ERK and P38, are key signaling proteins in MAPK pathway that are involved in the regulation of adipocyte proliferation, lipolytic function, and inflammation in adipose tissue (68–70), and they have been found to be modified by exercise training (71–73). The reduction in ERK phosphorylation we found after all three acute exercise sessions in our study suggests that a session of exercise may rapidly trigger signaling pathways that regulate the growth/differentiation of adipocytes (74). Because it is likely that ERK phosphorylation is essential for the early proliferative stages of adipogenesis in progenitor cells, but has to be shut-off for preadipocytes to continue to differentiate (68, 75), it is intriguing to speculate that the exercise-induced downregulation of ERK activity may elicit cell-type specific proliferative responses in progenitor cells and committed preadipocytes. In contrast, we did not observe difference in the phosphorylation of key lipolytic enzyme HSL or insulin signaling protein AKT, which aligns with our observation that the concentration of epinephrine, a lipolytic stimulator and insulin, an upstream activator of AKT returned to pre-exercise levels 1.5hours after exercise. Interestingly, despite the distinct differences in exercise intensity, duration, and energy expenditure among groups, the effect of acute exercise on the phosphorylation of the key metabolic proteins that we measured (ERK, P38, STAT3, HSL, AKT) was remarkably similar.

Our deconvolution analysis, which estimated cell-type proportions (%), revealed a post-exercise increase in macrophages and a reduction in adipocytes and vascular cells – expands our current understanding of the possibility of rapid cell-type-specific alterations in aSAT in response to acute exercise. While transcriptional alterations rather than changes in cell abundance could explain the observed shifts in cell-type proportions, our previous study found a reduction in a specific preadipocyte subgroup 12 hours after a 60-minute session of endurance exercise in individuals with obesity, suggesting that acute exercise may influence cell-type abundances rather quickly (16). Our finding of an increased proportion of macrophages after acute exercise aligns with a previous preclinical study that reported an increased abundance of anti-inflammatory macrophages in white adipose tissue 4 hours after acute swimming exercise in rats (76). While our data suggest that acute exercise may recruit macrophages and activate various immune cell types, it is important to note that inflammatory pathway upregulation in aSAT is largely short-lived (3), and long-term exercise training may even reduce the overall abundance of macrophages in aSAT (4). Although chronically elevated inflammation in adipose tissue and systemic circulation is commonly linked with a host of chronic diseases (77–79), paradoxically, acute inflammatory responses or immune cell activations in aSAT may be important contributors to favorable tissue remodeling (80). Whether aSAT cell-type proportions can be altered by exercise training is still unclear. However, our findings, along with previous findings from our lab (16) suggest that a session of endurance exercise may induce cell-type-specific responses that, when accumulated over time (i.e., training) may contribute to chronic adaptations in aSAT cellular proportions, as seen in mouse models (81).

Despite considerable efforts to control for various aspects of our study, some limitations remain. While participants served as their own control within each exercise group (i.e., pre/post exercise), the comparison across the three different exercise sessions was not conducted using a repeated measures design. Importantly, we were successful in tightly matching subjects in each of the three exercise groups for key factors that could impact the outcomes, such as, sex, cardiorespiratory fitness (VO_2_peak), body fat mass, and lean mass, enhancing confidence in the interpretations of our findings. Additionally, because we did not include a control trial in which adipose tissue samples were collected at the same time points without exercise, we cannot fully exclude the possibility that some of the observed exercise-induced responses in aSAT were influenced by factors unrelated to exercise, such as fasting duration or circadian rhythms. However, given the relatively short time between biopsies (<3h), it would be surprising if the robust changes we observed were solely driven by these factors. Furthermore, although our three different exercise treatments (LOW, MOD, and HIGH) represent commonly prescribed exercise programs, the three exercise sessions were not matched for exercise duration or energy expenditure, which may confound our interpretation. However, the goal of our study was to compare three distinct exercise modalities that are commonly implemented in clinical/applied settings, which makes matching for exercise duration and/or energy expenditure unfeasible. It is also important to acknowledge that the responses observed in the HIGH condition cannot be solely attributed to the high-intensity intervals themselves. The non-steady-state nature of this exercise may have influenced the observed adaptations as well. However, exercise intensity used during HIGH (which mimics common high-intensity interval training (HIIT) prescriptions) cannot be performed continuously for more than a few minutes at most. Another important limitation is that collecting only one aSAT sample ∼1.5 after exercise obviously does not fully capture the acute effect of exercise on aSAT that can persist for at least several hours or even a few days. Therefore, our findings are limited to characterizing the very early responses to exercise performed at low-, moderate- and high-intensity exercise.

In summary, our findings indicate that acute exercise can rapidly modify aSAT transcriptome, and some of these modifications were markedly different after the LOW, MOD, and HIGH exercise sessions in our study. Conversely, the three different exercise sessions evoked remarkably similar alterations in the expression of circadian genes, phosphorylation of some key metabolic proteins, and circulating cytokine concentrations, despite the vastly different exercise stimuli. Leveraging bioinformatic tools, we provide valuable insights about exercise-induced responses in aSAT, including the identification of gene groups that may be tightly linked with circulating cytokine concentrations during exercise and the possibility of changes in the proportions of different cell types in response to exercise. This work expands the understanding of molecular responses in aSAT triggered by a session of exercise that may contribute to the maintenance or remodeling of aSAT in regular exercisers, which in turn may be an important factor underlying the preserved cardiometabolic health often attributed to regular exercise.

## DATA AVAILABILITY

Differential analysis, pathway analysis, WGCNA module membership of genes are hosted on GitHub (https://github.com/ahnchi/3X-Study). All other datasets generated and analyzed in this study are available from the corresponding author upon reasonable request.

## CODE AVAILABILITY

R codes that were used for differential analysis and WGCNA are hosted on GitHub (https://github.com/ahnchi/3X-Study).

## ACKNOWLEDGMENTS

We are grateful to acknowledge the contributions of the study participants. We also acknowledge the excellent technical assistance from the Clinical Genomics Center at the Oklahoma Medical Research Foundation, and all the members of the Substrate Metabolism Laboratory. This study was supported by The National Institutes of Health (R01DK131724, P30DK089503, JFH). The funders had no role in study design, data collection and analysis, decision to publish or preparation of the manuscript.

## AUTHOR CONTRIBUTIONS

SCJP, CFB, and JFH designed the study. CA, TZ, GY, TR, OKC, SE, SJG, SM, RS, and JFH contributed to data acquisition. All authors have participated in data analysis and interpretation. CA and JFH drafted the work. All authors have participated in revising the work and approved the final version of the manuscript.

## COMPETING INTERESTS

The authors declare that they have no competing interests.

---

**Figure S1.**
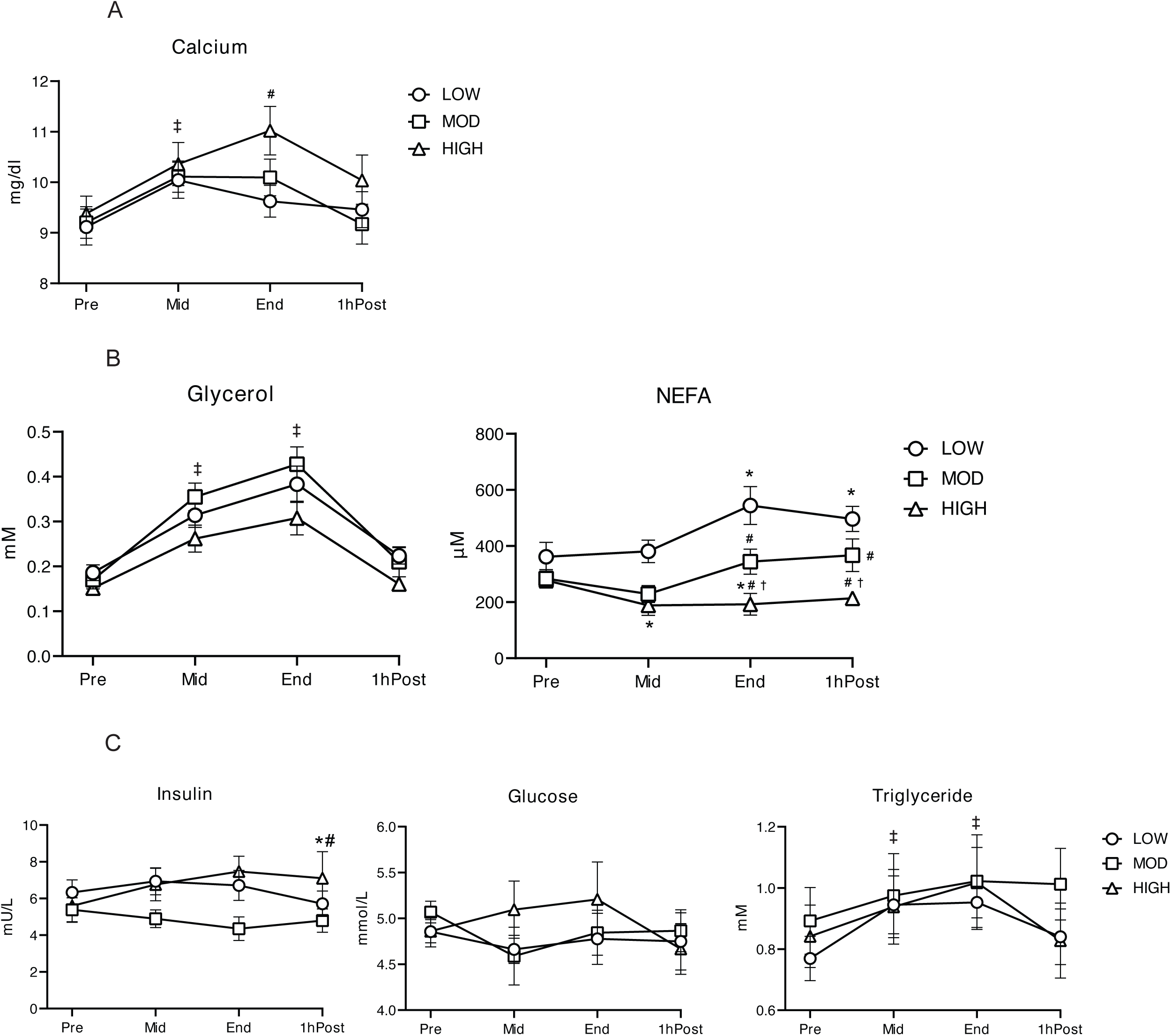
Concentrations of circulating calcium, glycerol, fatty acid, insulin, glucose, and triglyceride during and post LOW, MOD, and HIGH. A) Plasma calcium concentration. B) Plasma glycerol and fatty acid concentrations. C) Plasma insulin, glucose,and triglyceride concentrations. ‡main effect of time (p<0.05). *p<0.05 vs. Pre; †p<0.05 vs. MOD; #p<0.05 vs. LOW. Significant overall Time x Group interaction effects were detected in fatty acid (p<0.001, Type III ANOVA). There was a trend of Time (i.e., End) x Group (i.e., HIGH) interaction effect for calcium (p=0.06). Post-hoc pairwise comparison suggested significant difference in calcium concentration between LOW vs. HIGH at End (p=0.01). There was a significant Time (i.e., 1hPost) x Group (i.e., HIGH) interaction effect for insulin (p=0.025). Data is presented as Mean±SEM. Detailed p-values are included in the Result.

**Figure S2.**
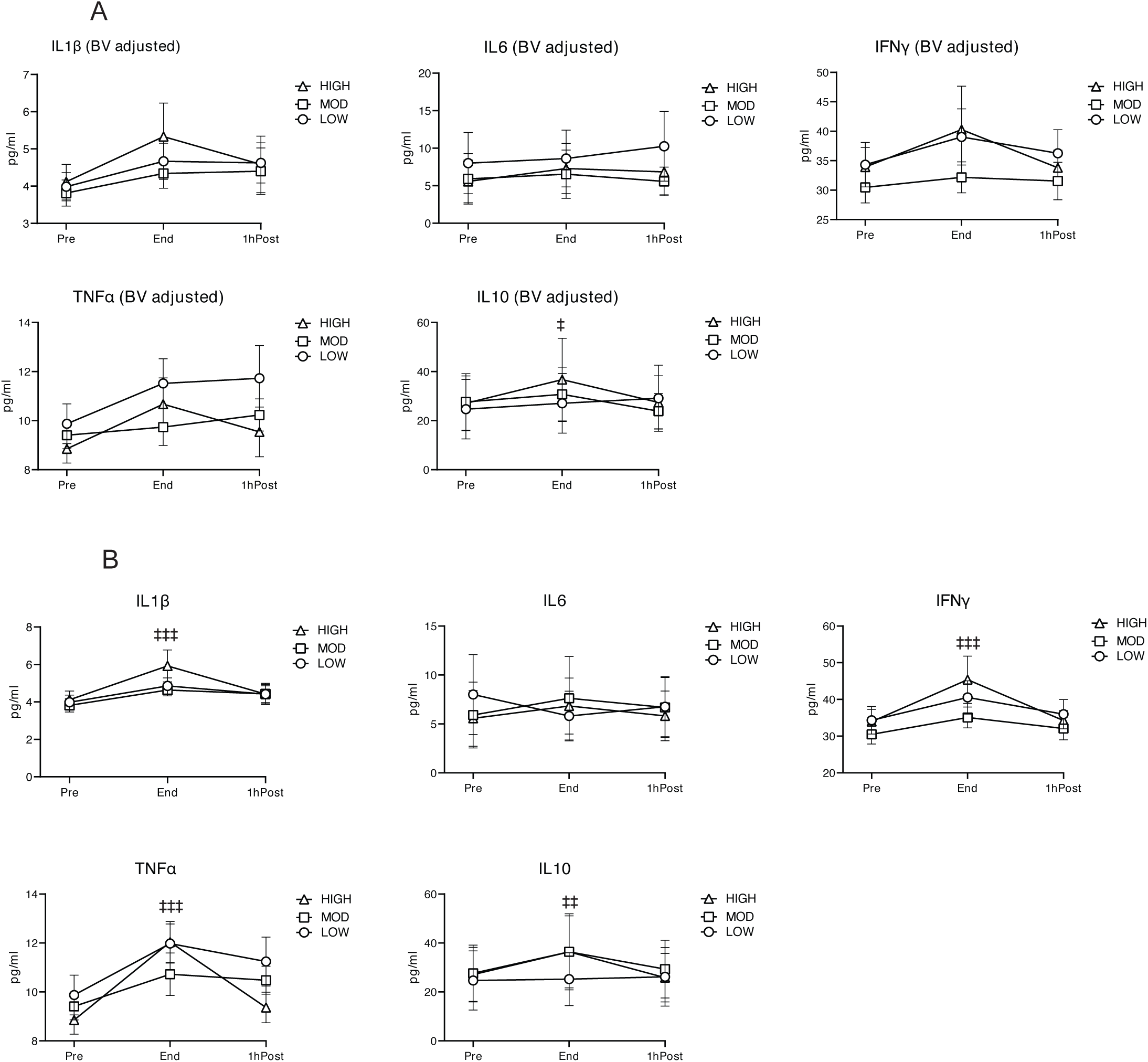
Concentrations of circulating cytokines during and post LOW, MOD, and HIGH. A) Blood volume-adjusted plasma IL1β, IL6, IFNγ,TNFα, IL10 concentrations. B) Unadjusted plasma IL1β, IL6, IFNγ,TNF α, IL10 concentrations. ‡main effect of time (p<0.05); ‡‡main effect of time (p<0.01); ‡‡‡main effect of time (p<0.001).. Sample sizes for IL10 – LOW: n=14; MOD: n=15; HIGH: n=14. Sample sizes for IL6 – LOW: n=10; MOD: n=14; HIGH: n=13. Sample sizes for IL1β, IFNγ, and TNFα - LOW: n=15; MOD: n=15; HIGH: n=15. Data is presented as Mean±SEM. BV, Blood volume.

**Figure S3.**
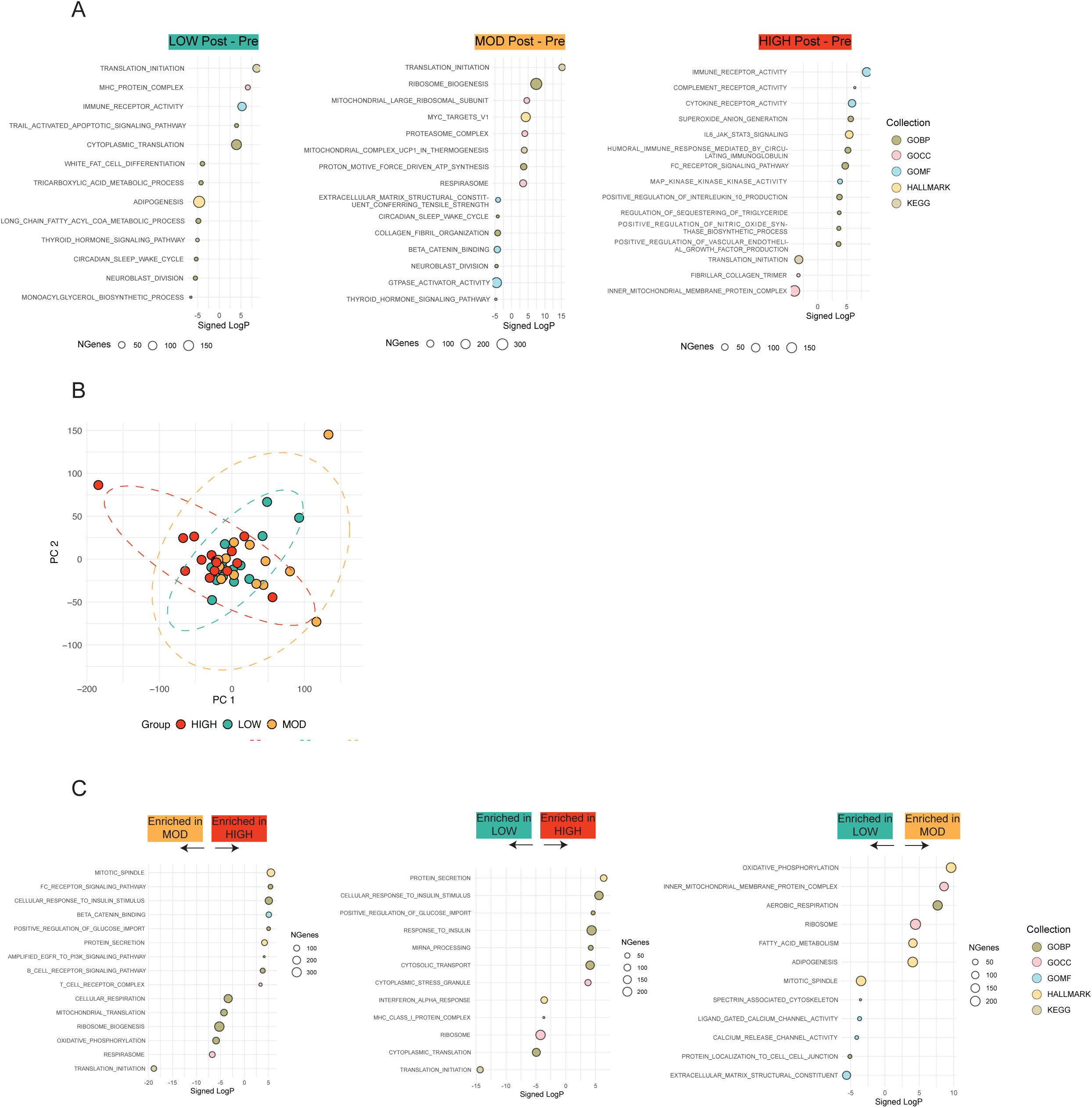
Gene set testing using CAMERA. A) Enriched biological pathways from each exercise treatment (comparing post- vs. pre-exercise), derived from CAMERA. Signed −log(p-value) from DESeq2 is used as the input for each gene. Positive values in the X-axis refers to upregulation after exercise. B) PCA of gene expression changes (delta of post/pre gene expression) in study subjects. Each point represents the subject. C) Enriched biological pathways from each exercise treatment (comparing exercise groups), derived from CAMERA. CAMERA, Competitive gene set test accounting for inter-gene correlation; PCA, Principal Component Analysis. Only significant terms (adjusted p<0.05) are shown in panel A and C.

**Figure S4.**
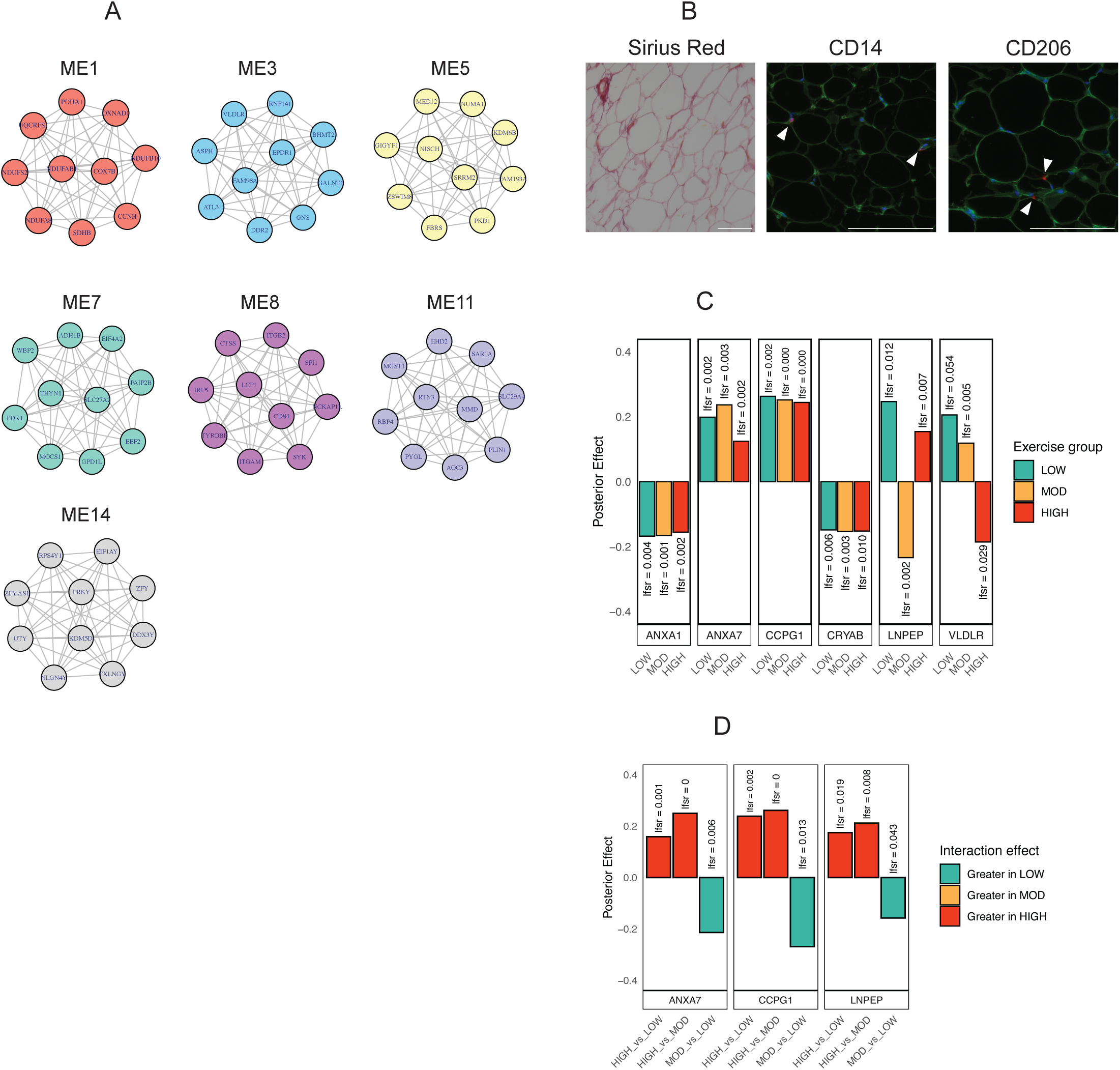
Integration of aSAT transcriptomics with clinical/tissue traits. A) Top 10 hub-genes from selected modules. Grey edges indicate intercorrelation between genes. B) Representative images of aSAT histology. Sirius Red was used to stain collagen type I and III depositions. CD14 was used as a marker for pro-inflammatory macrophages. CD206 was used as a marker for anti-inflammatory macrophages. Positive stains of CD14 and CD206 are marked with white arrows. C) DEGs among top 20 hub-genes of module 3. D) DEGs among top 20 hub-genes of module 3 that had significant group x interaction effect; ANXA7, CCPG1, and LNPEP.

## Notes

https://github.com/ahnchi/3X-Study

### Competing Interest Statement

The authors have declared no competing interest.

